# Mechanically-evoked spike responses of pentascolopidial chordotonal organs of *Drosophila* larvae

**DOI:** 10.1101/2023.05.26.542403

**Authors:** Ben Warren, Martin C. Göpfert

## Abstract

Stretch-sensitive ensembles of neurons in insects, known as Chordotonal organs (COs), function in proprioception, the detection of sound and substrate vibrations. Here we characterize the mechanical sensitivity of the lateral pentascolopidial CO (lch5) of *Drosophila* larvae to establish its postulated role in proprioception. We developed a physiologically realistic method to replicate proprioceptive input to lch5 by pulling the apodeme (tendon) to which the tips of the neurons attach. We found that lch5 sensory neurons respond transiently with a short latency to the velocity-component of stretch displacements and the release of stretch (relaxation). The mechanosensory mutant *inactive*, has a decreased response to mechanical stimuli and a lower overall spontaneous spike rate. Finally, we simulated the input that lch5 receives during crawling and observed spikes coincident with the start, middle and end of each peristaltic body contraction. We provide the first characterization of proprioceptive feedback in *Drosophila* larvae and firmly establish the proprioceptive function of lch5 in larval locomotion.

**Summary statement:** We have characterized the spiking responses of the pentascolopidial chordotonal organ of 20 Drosophila larvae to mechanical stimuli and reveal a clear role in proprioception.

## Introduction

Insects are equipped with an elaborate array of mechanoreceptors, essential for interacting with the environment and controlling body movements. Chordotonal organs (COs) are composed of bipolar stretch-sensitive neurons, which form elaborate frequency-discriminating acoustic detectors (Oldfield, 1982; Kamikouchi et al., 2009), function as proprioceptors in the context of locomotion (Usherwood et al., 1968), detect ground-borne vibration (Shaw, 1994) and serve a range of additional roles (Schnorbus, 1971; Field and Matheson, 1998). The pentameric CO of *Drosophila* larvae (lch5) has been shown to respond to sound and vibration (Scholz et al., 2015; Ohyama et al., 2015; Ohyama et al., 2013; Zhang et al., 2013) but characterization of lch5 responses to proprioceptive stimuli is lacking; despite its clear role in locomotion (Caldwell et al., 2003; Fushiki et al., 2013; Ohyama et al., 2013).

Proprioceptive COs, including the lch5 of *Drosophila* larvae, are typically formed of groups of scolopidia (a specialized ensemble of cells containing the mechanosensory neurons) which are attached between body segments via specialist attachment cells and an apodeme (tendon-like ligament). Well-studied examples include the metathoracic femoral CO (FCO), of the locust and stick insect which detect the relative angle between the tibia and femur. The mechanosensory neurons of the FCO alter their spike rate to either: a new resting position, during a change in resting position (velocity), acceleration, or a combination of these (Hofmann et al., 1985; Hofmann and Koch, 1985; Zill, 1985; 1989; Matheson, 1990; Büschges, 1994; DiCaprio et al., 2002). The neurons themselves, within any one CO, display heterogeneous responses to mechanical stimuli, known as range fractionation. For instance, any one position-sensitive neuron responds most strongly to a limited range of tibia-femur angles but together the FCO covers the entire range of angles (Hofmann et al., 1985; Zill, 1985). Similarly, any one velocity-sensitive neuron responds to a limited range of velocities but together the FCO codes for the full range of velocities (Matheson, 1992). Adding to the complexity of the FCO, individual, sensory neurons can be uni- or bi-directionally sensitive and respond to both an increase and decrease in tension. These heterogeneous response types in multiple COs provide detailed proprioceptive feedback necessary for coordination of locomotion.

Insect larvae possess a simplified array of COs essential to coordinate crawling (Caldwell et al., 2003; Fushiki et al., 2013; Titlow et al., 2014). Larvae CO-neurons are accessible for electrophysiological recordings (Zhang et al., 2013; Scholz et al., 2015) and fluorescent imaging (Ohyama et al., 2013; Ohyama et al., 2013); representing an attractive system to understand CO sensory input in locomotion. Our knowledge of the mechanical stimuli to which larvae CO respond is limited to responses to vibratory stimuli (Scholz et al., 2015; Ohyama et al., 2013; Ohyama et al., 2015; Zhang et al., 2013), which is thought to be a possible adaptation to detect predators (Zhang et al., 2013). Within each abdominal hemi-segment, of *Drosophila* larvae, are three singlet COs and one pentameric CO, lch5. The five bipolar neurons comprising lch5 is stretched diagonally from the dorsal posterior to the lateral anterior region of each abdominal segment; ideally placed to detect contractions necessary for crawling. It is generally assumed that, in addition to detecting vibration, lch5 responds to contractions of abdominal segments and that this sensory input is necessary for coordinating locomotion (Caldwell et al., 2003; Inbal et al., 2004; Fushiki et al., 2013; Ohyama et al., 2013).

Our first aim is to characterize the spiking response of lch5 to mechanical stimuli by recording extracellular spikes from the lch5 nerve. We delivered mechanical stimuli by stretching the cap cells onto which the tips of the neurons are embedded; this is a physiologically realistic way to simulate proprioceptive stimuli. Our second aim is to characterize the response of lch5 to proprioceptive displacements experienced during crawling to establish the role of lch5 in locomotion.

## Results

### Mechanical stimulation

We applied mechanical stimuli to lch5 by a piezo-actuator-coupled tungsten probe lowered onto the junction between the cap cells and cap attachment cells in lch5s in abdominal segments A2-A5 (Fig. 1A). Displacements were in-line with the long axis of lch5. We measured displacements of the probe and the cap cells, using the CCD camera imaged through differential interference contrast (DIC) optics (Fig. 1A). There was a linear Hookian relationship between probe and cap cell movements, though for larger displacements the cap cells moved relatively less, presumably as an elastic limit was reached (Fig. 1B). The range of displacements delivered to lch5 was limited to 15 μm which compares to the 462 ± 80 μm length of the lch5 from its two attachment sites. Movements beyond this resulted in the nerve of lch5 being pulled out of the recording electrode. Occasionally abdominal muscles contracted, as occurs in locomotion, and lch5 bent from straight to S-shaped. This constitutive contracted state rendered spike recordings from lch5 impossible. All recordings of lch5 were made in the relaxed non-contracted state.

**Figure 1.**
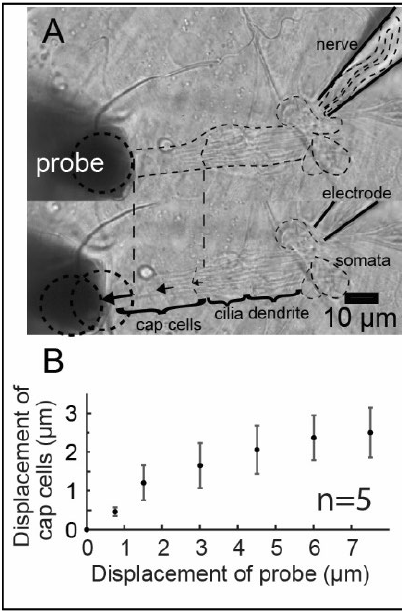
Mechanical stimulation of lch5 and relative displacement of probe and cap cells. Fig. 1 (A) Displacement of lch5 with tungsten probe visualized with DIC microscopy (B) Displacement of tungsten probe tip when *in situ* (when pressed against the attachment cells) against displacement of cap cells to step stimuli (n=5) ± SD.

### Spontaneous spiking properties of lch5

Spontaneous spikes were recorded at an average frequency of 46.6 ± 15.3 spikes per second (sp/s) (n=20) from the lch5 nerve. A spike-sorting algorithm was used to identify individual spike-forms (units) from lch5 (Fig. 2A). The spike frequency of distinguishable spike-forms had a range of 1.5 to 78.4 Hz with an average of 24.6 ± 23.2 Hz (n=34 neurons from 11 lch5s). The number of distinguishable spike-forms ranged from one to five, representing the five sensory neurons in lch5 (Fig. 2B). It is possible that spike-forms from multiple neurons were indistinguishable from the spike-sorting algorithm and that the number of neurons spontaneously spiking is underestimated from our analysis. However, the recruitment of spike forms responding to mechanical stimulation (see results section: *Velocity sensitive lch5 units*), suggests that, in the quiescent state, only a fraction of lch5 neurons were tonically spiking.

**Figure 2.**
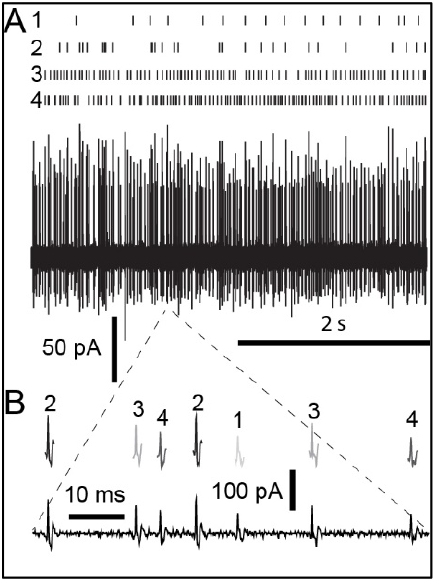
Extracellular recording of spontaneous spiking activity from lch5 and spike sorting. Fig. 2 (A) Extracellular currents recorded from lch5 nerve (middle trace), with raster plot of four spike types and (B) zoom in of time trace (bottom) showing the four spike types.

### Spike latency and response to step stimuli

In response to step displacements of 7.5 μm amplitude (away from lch5, defined as negative displacement), lch5 responded with spikes (Fig. 3A, B) within 2.36 ± 0.68 ms (n=10) (Fig. 3C). Spike rate significantly increased during the 10 ms after displacement (Fig. 3B) (LM: t=6.95_(20)_, p<4×10^−7^) comparing 10, 1 ms bins before and after displacement). These spikes were of similar amplitude and waveform of spontaneous spikes suggesting they were not artifacts (Fig. 3i, ii).

**Figure 3.**
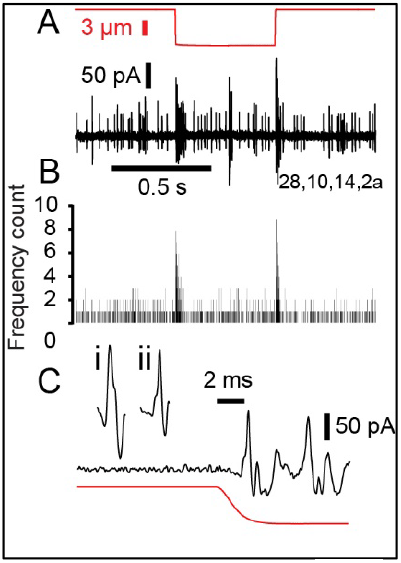
Latency of spikes recorded from lch5 in response to step displacements. Fig. 3 (A) Extracellular recording from lch5 nerve (black) in response to step displacement (red). (B) Histogram of seven consecutive recordings from the same lch5 with 1 ms bins. (C) Extracellular recording and step displacement on expanded time axis showing a step elicited spike (Ci) and a spontaneous spike (Cii).

### Lch5 neurons are velocity sensitive

To characterize proprioceptive-like responses of lch5 neurons we used ramp-and-hold displacements, in both directions (away from (negative) and toward (positive) lch5) with 1 s rise and fall times (Fig. 4Ai, ii). Ramp-and-hold displacements are an established method for identifying position-, velocity-, acceleration- and bi-directionally-sensitive proprioceptive neurons. Spike rate increased typically at ramp onset for both push and pull mechanical stimulation (Fig. 4Ai, ii). Some lch5s had a clear response confined to either the pull or push ramps (Fig. 4Aiii, asterix). To identify lch5s that responded to either positive or negative ramp stimulations, significant increases in spike rate were classified, if the number of spikes in the 100 ms bins (indicated with asterix), were at least three-times the standard deviation of the mean spike number of the remaining 100 ms bins. This analysis of lch5 responses to ramp-and-hold displacements yielded six lchs which responded to only positive or only negative ramp-and-hold displacements respectively, the histograms of which are plotted (Fig. 4Bi, ii). The proportion of 12 lch5s responding to each phase of the ramp and hold mechanical stimulus ranged from 9/12 for ramp onset for push stimulation down to 2/12 for ramp offset for push stimulation (Fig. 4Bi, ii).

**Figure 4.**
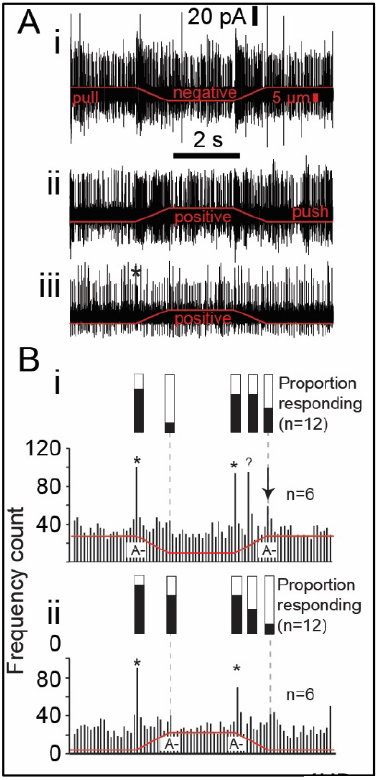
Extracellular spike responses of lch5 to ramp and hold displacements when pulling and pushing the apodeme. Fig. 4 (A) Extracellular recording from an lch5 nerve in response to a (Ai) pulling (negative) ramp and hold displacements and a pushing (Ai, Aii) (positive) direction. (B) Histogram of spike responses of six lch5 with 100 ms bins to negative (Bi) and positive (Bii) displacements. The proportion of 12 lch5s responding is indicted above by black infill to the bars.

A significant, increase in spike rate, occurred midway through the off ramp for negative displacements (Fig. 4B question mark) (LM: t_(76)_=6.84, p=9×10^−10^, comparing two 100ms bins of response with remaining 100 ms bins). Displacement-correlated responses were not limited to 100 ms following ramp *onset* of step-and-hold stimuli (Fig. 4B, arrow). There was a significant increase in spike rate with ramp *offset*, for negative displacements (Fig. 4B arrow) (LM: t_(76)_=2.45, p=0.008,, comparing two 100ms bins of response with remaining 100 ms bins).

### Velocity/acceleration threshold of Lch5 neurons

We stretched lch5 at different velocities but with the same displacement (Fig. 5A). This protocol determined the threshold velocity/acceleration necessary to elicit a significant increase in overall neuronal spiking. Histograms with 5 ms bins were generated from spiking responses. When the number of spikes per bin was more than six-times above the standard deviation (i.e. significantly above the spontaneous spiking rate) of all bins over the four second recording period, we classified this as a response above threshold (Fig. 5B). There was considerable spread of velocity and acceleration thresholds from 5-45 μm/s and 1.6-11.2 μm/s^2^ (Fig. 5C). The increase in spiking rate was transient in response to the velocity/acceleration component of mechanical stimuli, i.e. there was no tonic spiking. Such transient responses at the beginning of a velocity stimulus are also indicative of acceleration-sensitive units, although acceleration-sensitive units responses are usually limited to only one spike in femoral chordotonal organs (Matheson, 1990; Hofmann and Koch, 1985; Büschges, 1994).

**Figure 5.**
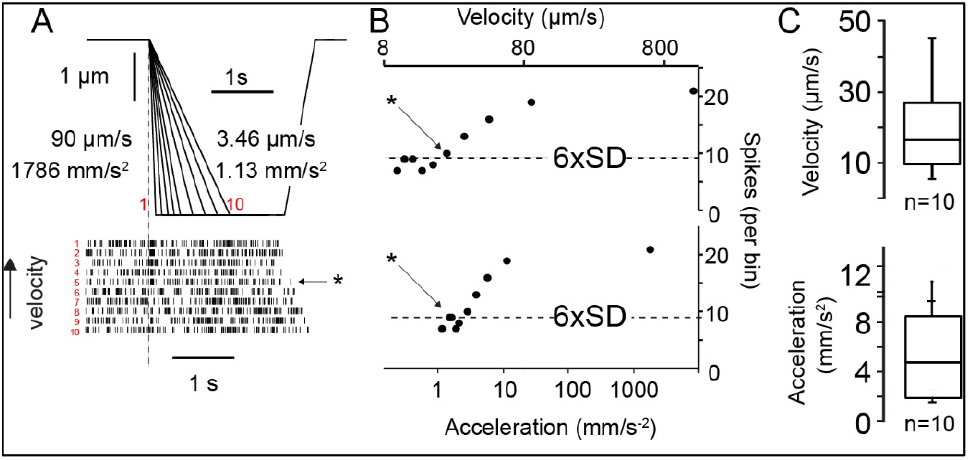
Experimental protocol to determine the velocity/acceleration threshold of lch5. Fig 5 (A) Displacement of tungsten probe and raster plot of lch5 spikes to different velocities and accelerations of displacement (90 μm/s^1^ to 3.46 μm/s^1^ and 1786 mm/s^2^ to 1.13 mm/s^2^). (B) Plot displaying the spikes per 5 ms bin in response to different ramp velocities and accelerations, asterisk (also in (A)) corresponds to threshold velocity/acceleration, when the number of spikes is above six times the standard deviation (i.e. significantly above noise). (C) Quantification of the velocity and acceleration thresholds found for ten separate lch5s.

**Figure 6.**
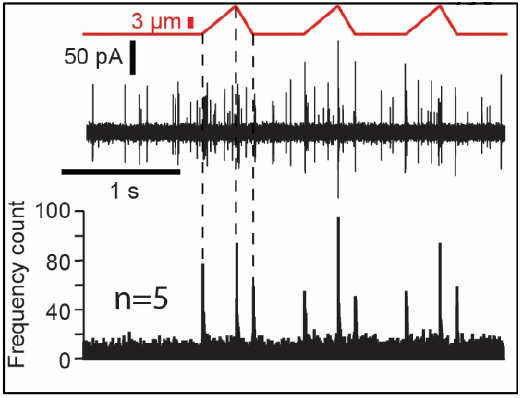
Spike responses to simulated peristaltic contractions. Fig. 7 Spike responses of lch5 (middle) to displacements mimicking three cycles of peristaltic contractions (top) and histogram of spike responses using 2.5 ms bins (bottom) of 10-20 repetitions in five lch5.

### Lch5 responds to the dynamic components of crawling

To provide physiologically realistic proprioceptive input to lch5 we used a pattern of probe movement that mimicked movements measured in the muscles of *Drosophila* larvae (Heckscher et al., 2012) (Fig. 7A, red trace). We repeated the crawling stimulus every one second as this is a typical period for *Drosophila* larvae crawling contractions (Wang et al., 1997; Saraswati et al., 2003). The lch5 neurons respond transiently and consistently to the onset and offset of movement. These three phases of the cycle correspond to the 1) start of muscle elongation 2) change to muscle contraction and 3) when muscles stop contractions (Fig. 7).

### Inactive mutant phenotypes

We investigated the spiking properties of lch5 in the mutant *inactive (iav)*. Inactive forms a heteromeric complex with Nanchung and its loss severely affects the mechanical properties of the adult fly’s Johnston’s (chordotonal) organ and sound-evoked compound potentials recorded from JO neurons (Gong et al., 2004; Göpfert et al., 2006). lch5 of *iav*^*1*^ null mutants exhibited a greatly reduced spontaneous firing rate (4.2 ± 7.2 Hz, n=11) compared to Canton S wild-type controls (46.6 ± 15.3 Hz, n=20) (Fig. 8A). The responses of *iav*^*1*^ mutants to 20 Hz sinusoidal stimulation were also significantly attenuated across a range of displacements (Fig. 8B).

**Figure 8.**
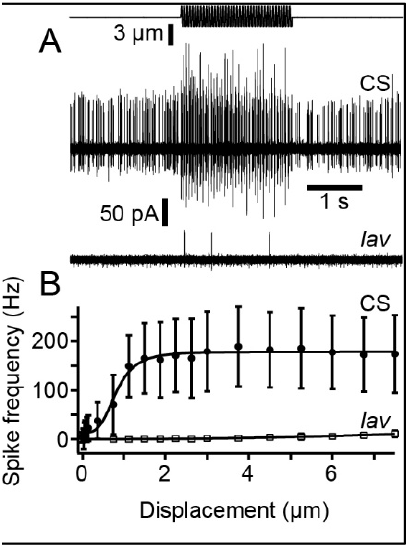
Response of *inactive* and Canton S control to 20 Hz sinusoidal stimulation of different amplitudes. Fig. 8 (A) Stimulus displacement (top) and extracellular spike responses of Canton S (CS) and *Iav*^*1*^. (B) Quantification of spike frequency increase ± SD to different amplitudes of Canton S (n=12) and *Iav*^*1*^ (n=11).

## Discussion

The genetic amenability of *Drosophila* has made it an excellent model organism, however its tiny size has hampered efforts to investigate electrophysiological properties of individual sensory cells in COs. In contrast, intracellular recordings have been performed in many proprioceptive COs in other insects, which lack the genetic tools available in *Drosophila* to generate Ch-specific mutations (Hofmann et al., 1985; Zill, 1985; Field and Pflüger 1989; Matheson 1990; Büschges 1994). Extracellular recordings from COs in *Drosophila* larvae have previously been performed (Zhang et al., 2013; Ohyama et al., 2015; Scholz et al., 2015) but investigation of their physiology has been limited to responses to vibrational stimuli. This study investigates the response of lch5 to proprioceptive lower frequency stimuli using extracellular recordings and spike detection. We measured quantitative differences in spike rate and spike response between Canton S controls and the *iav*^*1*^ mutants.

Our findings support a simple viscoelastic apodeme in Drosophila larvae with an elastic limit (Fig. 1B). The anatomy of CO apodemes in adult insects are more complex (Shelton et al., 1992; Nowel et al., 1995). For instance, the apodeme of the metathoracic FCO of the locust has multiple ligaments which are sequentially stretched and recruited and could give rise to range fractionation (Field 1991; Shelton et al., 1992). In lch5 of *Drosophila* larvae all cap cells attach to the same point and when inspected through DIC microscopy have homogenous appearance.

In the absence of mechanical stimulation individual sensory neurons of lch5 spontaneously spike at a rate of 24.6 ± 23.2 Hz. Spontaneous spiking is a feature typical of proprioceptive COs in insects which respond to mechanical stimulation by altering their spike rate above or below their spontaneous rate (Hofmann et al., 1985; Hofmann and Koch, 1985; Zill, 1985; Field and Pflüger, 1989; Matheson, 1990; Büschges, 1994; DiCaprio et al., 2002). The lateral pentascolopidial CO (lch5) in *Drosophila* larvae responds to mechanical step stimuli with a short-latency change in the spiking rate 2.36 ± 0.68 ms (n=11). This latency is similar to acceleration-sensitive receptors in the stick insect FCO (Hofmann and Koch, 1985) but lies above that measured in acoustic responses of the Johnston’s organ of *Drosophila* (∼500 μs latency) (Albert et al., 2007). Latencies here could be limited by the time taken for the apodeme to stretch to a threshold amount. Our measurements show that the long, slender attachment cells of lch5 behave as a viscoelastic material but sudden movements, such as step stimuli used here, may cause a delay in stretching of the apodeme as viscous effects momentarily overcome elasticity.

The sensory neurons of lch5 fire transiently in response to a change in displacement. The sensory neurons of lch5 are either responding transiently to velocity, with an average threshold of 19.1 ± 7.2 μm/s, or are acceleration-sensitive, with an average threshold of 4.61 ± 2.9 μm/s^2^. Either way the transient-phasic nature of these responses indicates that lch5 senses and provides feedback for dynamic movements and that the distance between body segments is not signaled by lch5. The other main type of sensory neuron, multidendritic neurons that completely tile the body wall, seem well placed to provide feedback on the longitudinal state of body segments (Grueber et al., 2002). In particular type I multidendritic and bipolar neurons, the dendrites of which span the width and length of each segment, appear to provide most of the sensory feedback necessary for normal locomotion (Hughes and Thomas, 2007). Their presumed ability to signal length in body segments may make position-sensitive CO sensory neuron responses redundant in soft-bodied larvae; an idea first suggested by Hughes and Thomas (2007).

Although peristaltic contractions necessary for larval locomotion are still present when sensory input is disrupted, sensory feedback is necessary for altering central pattern generator output and for normal locomotive behaviour (Caldwell et al., 2003; Hughes and Thomas, 2007; Song et al., 2007; Fushiki et al., 2013; Titlow et al., 2014;). A goal in the field is to identify the sensory neurons necessary and sufficient for providing sensory feedback for locomotion. The relative role of CO neurons and multidendritic neurons remains contested. Temperature-sensitive *Shibire* expression to impair each neuron type resulted in different conclusions on crawling speed. Whilst two studies found a decrease in crawling speed when only multidendritic neurons were inhibited (Hughes and Thomas, 2007; Song et al., 2007), another found that larval crawling speed was decreased by inhibiting only COs (Fushiki et al., 2013). A further study using CO mutants found a decreased speed of crawling compared to controls (Caldwell et al, 2003).

Hughes and Thomas (2007) have suggested a “mission accomplished” model of sensory feedback based on inferences of how multidendritic neurons operate in adult insects. Bipolar and class I multidendritic neurons provide the “mission accomplished” signal by an increase or decrease of their spike rate. This signal is necessary in order to coordinate the contraction and relaxation of muscles in the local and adjacent segments. Our simulation of peristaltic contractions delivered to lch5 revealed that a discrete number of spikes closely follow the start middle and end of a simulated body contraction (Fig. 7). Thus signals from lch5 could provide a “mission accomplished” signal, at the end of each three phases of a crawling cycle which would be integrated with sensory input from multidendritic neurons to influence motor output.

The COs in *Drosophila* larvae have been shown with calcium imaging and focal extracellular recordings to be sensitive to vibration (Ohyama et al., 2015; Ohyama et al., 2013; Zhang et al., 2013; Scholz et al., 2015). Detection of sound is thought to be important for *Drosophila* larvae to avoid predators (Zhang et al., 2013). We were unable to elicit responses of lch5 to sound of various frequencies from a loud speaker, even at high sound pressure levels > 100 dB SPL, which suggests that the lch5 responses are fundamentally vibration-(and not sound-) sensitive. A parallel proprioceptive role of larval COs in *Drosophila* is also suggested by the study of locomotion phenotypes where COs are genetically compromised (Caldwell et al., 2003; Wu et al., 2011; Zhou et al., 2012; Fushiki et al., 2013; Zhang et al., 2013), and also because lch5 spans across body segments (Klein et al., 2010). Proprioceptive FCOs in insects that have evolved into dedicated vibration receptors, for example the subgenual organ of the cockroach, bear morphological specializations, whereby it no longer spans adjacent body segments (Schnorbus, 1971; Moran and Rowley, 1975; Shaw, 1994). This study provides *in vivo* extracellular recordings of lch5 showing responses to simulated peristaltic contractions. The lch5 appears to serve a dual sensory role; responding to vibration and proprioceptive cues. To investigate the extent to which lch5 is a vibration/proprioceptive sensory organ, both types of stimuli need to be delivered in the same experimental setup. We were unable to investigate vibrational responses here as our dedicated proprioceptive setup was unable to stimulate lch5 at frequencies over 100 Hz.

To date, investigations of *Drosophila* larval locomotion have used the powerful genetic tools afforded in *Drosophila* combined with behavioural analysis. Systematic physiological analysis of sensory neurons in response to proprioceptive stimuli is now needed to further identify the sensory neurons necessary for proprioceptive feedback and to start to build realistic models. Such multimodal analysis certainly seems warranted as signals from different sensory neuron types are well integrated and converge throughout the integrated nervous system of *Drosophila* larvae (Ohyama et al., 2015). Quantitative differences in sensory neuron function of mutations affecting COs in larvae have been shown with calcium imaging (Zhang et al., 2013). We now add another level of analysis by finding quantitative differences in the spontaneous spike rate and spike response to sinusoidal stimuli of sensory neurons of *iav*^*1*^ mutants when compared to control larvae. Such findings set the stage for using the ‘simpler’ locomotory system of *Drosophila* larvae, to understand how COs contribute to proprioceptive input.

## Method

### Preparation

Third instar larval *Drosophila melanogaster* (Miegen, 1830), Insecta, Diptera, Drosophilidae] were pinned to an agar coated recording chamber. Larvae were prepared using a modified fillet preparation in haemolymph-like saline containing in mM 103 NaCl, 3 KCl, 5 2-([1,3-dihydroxy-2-(hydroxymethyl)propan-2-yl]amino)ethanesulfonic acid, 10 trehalose, 10 glucose, 7 sucrose, 26 NaHCO_3_, 2CaCl_2_, 1 NaH_2_PO_4_, 4 MgCl_2_, adjusted to 7.2 with KOH. For the fillet preparation two pins (0.1 × 1 mm tungsten wire) were inserted through the mouth hooks and tail, pinning the larva dorsal side up before saline was added. We used ultra-fine clipper scissors (Model #15300-00, Fine Science Tools, Heidelberg, Baden-Württemberg, Germany) to cut along the dorsal midline, and a further four pins were used either side of the mouth hooks and tail pins to pin down the corners of the cuticle. All internal organs, body fat and two main trachea were removed with fine forceps but the muscles and nervous system were left intact.

### Piezo stimulation and electrophysiology

A piezo-actuator and controller (Physik Instrumente GmbH, Karlsruhe, Baden-Württemberg, Germany) (P-841.10, E-709.SRG) under the control of Patchmaster software (HEKA, Lambrecht, Niedersachsen, Germany) was used to control a custom-made tungsten probe (18 mm length, 0.2 mm diameter), on top of the tendon (apodeme) of lch5, ∼40 μm distal to the cap cells at the point where the tendon is narrowest. The tungsten probe was inline with the orientation of the attachment and ligament cells. Suction pipettes made from borosilicate glass (Science Products GmbH, Hofheim, Hesse, Germany) (GB150T-8P) pulled with a P-1000 (Sutter instrument, Novato, California, USA) electrode puller to a tip diameter ∼4 μm and filled with saline described above, were used to suck the lch5 nerve (∼20 □m) into the pipette. After this, gentle suction was used to attach the pipette to the sheath surrounding the somata of lch5. A HEKA patch-clamp amplifier (EPC 10 USB) under the control of Patchmaster software (HEKA) was used in voltage-clamp mode and data were filtered online with an analog 4-pole low-pass Bessel filter at 2.9 kHz. Data were sampled at 20 kHz.

### Microscopy, CCD imaging and Doppler laser recordings

An Examiner D.1 microscope (Zeiss, Oberkochen, Baden-Württemberg, Germany) with a 63X objective (Zeiss, 424516-9041) and DIC slider (Zeiss, 426961) and an AxioCam MRm (Zeiss, 1388×1040 pixels) under the control of Axiovision (Zeiss V.4.8.2.0) were used to image the probe and neuron displacements. A Doppler laser (Polytec, PSV-400, Harpenden, Hertfordshire, UK) was used to confirm displacements of the probe. Sinusoidal stimuli delivered to the probe delivered reliable amplitude up to between 60 and 100 Hz. We used a white noise stimulus to displace the tungsten probe and record its frequency response. The tungsten probe exhibited an almost flat frequency response within the frequencies used (Fig. 1A).

### Inactive mutants

Canton S larvae were used as wild-type strains. *Iav*^1^ mutants as described previously (Gong et al., 2004) were obtained from the Bloomington Stock Center.

### Data analysis and Statistics

Igor software (Wavemetrics, Lake Oswego, Oregon, USA) (version 6.3.2.3) was used to high-pass filter the data (Finite Impulse Response Filter, end of reject band, start of pass band 500, 555 Hz). Spike 2 software (Cambridge Electronic Design, Cambridgeshire, UK) (version 7) was used for spike sorting. The time window was set to 0.4 ms before and after the peak amplitude of spikes. The threshold of spike detection was 1.8 pA. Automatic template generation was used and spikes were classified as the same when within 10% of the same amplitude. Statistical tests were computed in excel and all error bars are standard deviation. Linear Models (LM) were used for pairwise comparisons.

## Abbreviations

CO: Chordotonal Organ
FCO: Femoral chordotonal Organ
lch5: lateral pentameric chordotonal organ

## Author Contributions

B.W. Designed experiments, collected and analysed data and wrote the manuscript. M. C. G. designed experiments and wrote the manuscript.

## Acknowledgements

We would like to thank Heribert Gras for advice on experimental protocols and Tom Matheson for comments on the manuscript.

## References

Albert JT, Nadrowski B, Göpfert MC (2007). Mechanical signatures of transducer gating in the Drosophila ear. Curr. Biol. 17, 1000–1006. DOI: 10.1016/j.cub.2007.05.004

Adams CM, Anderson MG, Motto DG, Price MP, Johnson WA, and Welsh MJ (1998) Ripped Pocket and Pickpocket, Novel Drosophila DEG/ENaC Subunits Expressed in Early Development and in Mechanosensory Neurons J. Cell Biol. 140:143–152. DOI: 10.1083/jcb.140.1.143

Bharadwaj R, Roy M, Ohyama T, Sivan-Loukianova E, Delannoy M, Lloyd TE, Zlatic M, Eberl DF and Kolodki AL (2011). Cbl-associated protein regulates assembly and function of two tension-sensing structures in Drosophila. Development 140, 627–638. DOI:10.1243/dev.0855100

Büschges A. (1994). The physiology of sensory cells in the ventral scoloparium of the stick insect femoral chordotonal organ. J. Exp. Biol. 189, 285–292. DOI:10.1242/jeb.189.1.285

Caldwell JC, Miller MM, Wing S, Soll SR and Eberl DF (2003). Dynamic analysis of larval locomotion in Drosophila chordotonal organ mutants. Proc. Natl. Acad. Sci. USA 100, 16053–16058. DOI: 10.1073/pnas.2535546100

DiCaprio RA, Wolf H and Büschges A (2002). Activity-dependent sensitivity of proprioceptive sensory neurons in the stick insect femoral chordotonal organ. J. Neurophysiol.88, 2387–2398. DOI: 10.1152/jn.00339.2002

Field LH and Pflüger H-J (1989). The femoral chordotonal organ: A bifunctional orthopteran (Locusta migratoria) sense organ? Comp. Biochem. Physiol. 93, 729–743. DOI: 10.1016/0300-9629(89)90494-5

Field LH (1991) Mechanism for range fractionation in chordotonal organs of Locusta migratoria (L) and Valanga sp. (Orthoptera: Acrididae) Int. J. Insect Morphol. Embryol. 20, 25–39. DOI: 10.1016/0020-7322(91)90025-5

Field LH. and Matheson T (1998). In Advances in insect physiology 27: Chordotonal organs of insects (ed. P. Evans and V. Wigglesworth), pp. 1–230. London: Academic Press Ltd. DOI: 10.1016/S0065-2806(08)60013-2

Fushiki A, Kohsaka H and Nose A (2013). Role of sensory experience in functional development of drosophila motor circuits. PLoS ONE. 8, e62199. DOI: 10.1371/journal.pone.0062199

Gong Z, Son W, Chung YD, Kim J, Shin DW, McClung CA, Lee Y, Lee HW, Chang DJ, Kaang BK, Cho H, Oh U, Hirsh J, Kernan MJ and Kim C. (2004). Two interdependent TRPV channel subunits, Inactive and Nanchung, mediate hearing in Drosophila J. Neurosci. 24, 9059–9066. DOI: 10.1523/JNEUROSCI.1645-04.2004

Göpfert MC, Briegel H and Robert D (1999). Mosquito hearing: sound-induced antennal vibrations in male and female Aedes Aegypti. J. Exp. Biol. 202, 2727–2738. DOI: 10.1242/jeb.2020.20.2727

Göpfert MC, Albert JT, Nadrowski B and Kamikouchi A (2006). Specification of auditory sensitivity by Drosophila TRP channels. Nat. Neurosci. 9, 999–1000. DOI: 10.1038/nn1735

Grueber WB, Jan LY and Jan YN (2002). Tiling of the drosophila epidermis by multidendritic sensory neurons. Development 129, 2867–2878. DOI: 10.1242/dev.129.12.2867

Heckscher ES, Lockery SR and Doe CQ (2012). Characterization of Drosophila larval crawling at the level of the organism, segment, and somatic body wall musculature. J. Neurosci. 32, 12460–12471. DOI: 10.1523/JNEUROSCI.0222-12.2012

Hofmann T and Koch UT (1985). Acceleration receptors in the femoral chordotonal organ of the stick insect, Cuniculina impigra. J. Exp. Biol. 114, 225–237. DOI: 10.1242/jeb.114.1.225

Hofmann T, Koch UT and Bässler U (1985). Physiology of the femoral chordotonal organ in the stick insect, Cuniculina impigra. J. Exp. Biol. 114, 207–223. DOI: 10.1242/jeb.114.1.207

Hughes CL and Thomas JB (2007) A sensory feedback circuit coordinates muscle activity in Drosophila. Mol. Cell Neurosci. 35, 383–396. DOI: 10.1016/j.mcn.2007.04.001

Hustert R (1982). The proprioceptive function of a complex chordotonal organ associated with the mesothoracic coxa in locusts. J. comp. Physiol. A. 147, 389–399. DOI: 10.1007/BF00609673

Inbal A, Volk T, Salzberg A (2004) Recruitment of ectodermal attachment cells via an EGFR-dependent mechanism during the organogenesis of Drosophila proprioceptors. Dev. Cell 7, 241–250. DOI: 10.1016/j.devcel.2004.07.001

Kamikouchi A, Inagaki HK, Effertz T, Hendrich O, Fiala A, Göpfert MC and Ito K (2009). The neural basis of Drosophila gravity-sensing and hearing. Nature 458, 165–171. DOI: 10.1038/nature07810

Kaplin JM, Horvitz HR (1993) A dual mechanosensory and chemosensory neuron in Caenorhabditis elegans. PNAS 90, 2227–2231. DOI: 10.1073/pnas.90.6.2227

Klein Y, Halachmi N, Egoz-Matia N, Toder M, Salzberg A (2010). The proprioceptive and contractile systems in Drosophila are both pattered by the EGR family transcription factor stripe. Dev. Biol. 337, 458–470. DOI: 10.1016/j.ydbio.2009.11.022

Lee Y, Lee Y, Lee J, Bang S, Hyun S, Kang J, Hong ST, Bae E, Kaang BK, Kim J (2005) Pyrexia is a new thermal transient receptor potential channel endowing tolerance to high temperatures in Drosophila melanogaster Nat. Genet. 37, 305–310. DOI: 10.1038/ng1513

Lehnert BP, Baker AE, Gaudry Q, Chiang A-S, Wilson RI (2013). Distinct roles of TRP channels in auditory transduction and amplification in Drosophila. Neuron 77, 115–128. DOI: 10.1016/j.neuron.2012.11.030

Marley R. and Baines R. A. (2011). Dissection of first- and second-instar Drosophila larvae for electrophysiological recording from neurons: the flat (or fillet) preparation. Cold Spring Harb. Protoc. Sep 1, 9. DOI: 10.1101/pdb.prot065649

Marshall KL and Lumpkin EA (2012). The molecular basis of mechanosensory transduction. Adv. Exp. Med. Biol. 739, 142–155. DOI: 10.1007/978-1-4614-1704-0_9

Matheson T (1990). Responses and locations of neurons in the locust metathoracic femoral chordotonal organ. J. Comp. Physiol. A 166, 915–927 DOI: 10.1007/BF00187338

Matheson T (1992). Range fractionation in the locust metathoracic femoral chordotonal organ J. Comp. Physiol. A. 170, 509–520. DOI: 10.1007/BF00191466

Moran DT and Rowley JC (1975). The fine structure of the cockroach subgenual organ. Tissue and Cell 7, 91–106. DOI: 10.1016/S0040-8166(75)80009-7

Nowel MS, Shelton PMJ and Stephen RO (1995). Functional organization of the metathoracic chordotonal organ in the cricket Acheta domesticus. J. Exp. Biol. 198, 1977–1988. DOI: 10.1242/jeb.198.9.1977

Ohyama T, Schneider-Mizell CM, Fetter RD, Aleman JV, Franconville R, Rivera-Alba M, Mensh BD, Branson KM, Simpson JH, Truman JW, Cardona A, Zlatic M (2015). A multilevel multimodal circuit enhances action selection in Drosophila. Nature 520, 633–639. DOI: 10.1038/nature14297

Ohyama T, Jovanic T, Denisov G, Dang TC, Hoffmann D, Kerr RA and Zlatic M (2013). High-throughput analysis of stimulus-evoked behaviours in Drosophila larvae reveals multiple modality-specific escape strategies. PLoS ONE 8, e71706. DOI: 10.1371/journal.pone.0071706

Oldfield BP (1982). Tonotopic organisation of auditory receptors in Tettigoniidae (Orthoptera: Ensifrea). J. Comp. Physiol. A 147, 461–469. DOI: 10.1007/BF00612011

Saraswati S, Fox LE, Soll DR and Wu C-F (2004). Tyramine and octopamine have opposite effects on the locomotion of drosophila larvae. J. Neurobiol. 58, 425–441. DOI: 10.1073/pnas.1813554116

Schnorbus H. (1971). Die subgenualen sinnesorgane von Periplaneta Americana: histologie und vibrationsschwellen. Z. vergl. Physiol. 71, 14–48. DOI: 10.1007/BF03395969

Scholz N, Gehring J, Guan C, Ljaschenko D, Fischer R, Lakshmanan V, Kittel RJ, Langenhan T (2015). The adhension GPCR latrophilin/CIRL shapes mechanosensation. Cell 11, 1–9. DOI: 10.1016/j.celrep.2015.04.008

Shaw SR (1994). Detection of airborne sound by a cockroach ‘vibration detector’: a possible missing link in insect auditory evolution. J. Exp. Biol. 193, 13–47. DOI: 10.1242/jeb.193.1.13

Shelton PMJ, Stephen RO, Scott JJA and Tindall AR (1992). The apodeme complex of the femoral chordotonal organ in the metathoaracic leg of the locust Schistocera gregaria. J. Exp. Biol. 163, 345–358. DOI: 10.1242/jeb.163.1.345

Schnorbus H (1971) Die subgenualen sinnesorgane von Periplaneta americana: Histologie und Vibrationsschwellen. Z. verla. Physiologie 71, 14–48. DOI: 10.1007/BF03395969

Sokabe T, Tsujiuchi S, Kadowaki T, Tominaga M (2008) Drosophila painless is a Ca2+-requiring channel activated by noxious heat. J. Neurosci. 28, 9929–9938. DOI:10.1523/JNEUROSCI.2757-08.2008

Song W, Onishi M, Jan LY, and Jan YN (2007). Peripheral multidendritic sensory neurons are necessary for rhythmic locomotion behavior in Drosophila larvae. Proc. Nat. Acad. Sci. 104, 5199–5204. DOI: 10.1073/pnas.0700895104

Stein W and Sauer AE (1999) Physiology of vibration sensitive afferents in the femoral chordotonal organ of the stick insect. J. Comp. Physiol. A 184, 253–263. DOI: 10.1007/s003590050323

Titlow JS, Rice J, Majeed ZR, Holsopple E, Biecker S and Cooper RL (2014). Anatomical and genotype-specific mechanosensory responses in Drosophila melanogaster larvae. Neurosci. Res. 83, 54–63. DOI: 10.1016/j.neures.2014.04.003

Tracey WD, Wilson RI, Laurent G, Benzer S (2003). painless, a Drosophila gene essential for nociception. Cell. 113, 261–273. DOI: 10.1016/s0092-8674(03)00272-1

Usherwood PNR, Runion HI and Campbell JI (1968). Structure and physiology of a chordotonal organ in the locust leg. J. Exp. Biol. 48, 305–323. DOI: 10.1016/0022-1910(74)90236-4

Wang JW, Sylwester AW, Reed D, Wu DAJ, Soll DR and Wu CF (1997). Morphometric description of the wandering behavior in Drosophila larvae: aberrant locomotion in Na+ and K+ channel mutants revealed by computer-assisted motion analysis. J. Neurogenetics 11, 231–254. DOI: 10.3109/01677069709115098

Wu Z, Sweeny LB, Ayoob JC, Chak K, Andreone BJ, Ohyama T, Kerr R, Luo L, Zaltic M and Kolodkin AL (2011). A combinatorial semaphoring code instructs the initial steps of sensory circuit assembly in the drosophila CNS. Neuron 70, 281–298. DOI: 10.1016/j.neuron.2011.02.050

Zhang W, Yan Z, Jan LY and Jan YN (2013). Sound response mediated by the TRP channels NOMPC, NANCHUNG, and INACTIVE in chordotonal organs of Drosophila larvae. Proc. Natl. Acad. Sci. USA 110, 13612–13617. DOI: 10.1073/pnas.1312477110

Zhong L, Hwang RY, Tracey WD (2010) Pickpocket is a DEG/ENaC protein required for mechanical nociception in Drosophila larvae. Curr. Biol. 20, 429–434. DOI: 10.1016/j.cub.2009.12.057

Zhong L, Bellemer A, Yan H, Honjo K, Robertson J, Hwang RY, Pitt GS, and Tracey WD (2012) Thermosensory and non-thermosensory isoforms of Drosophila melanogaster TRPA1 reveal heat sensor domains of a thermoTRP channel. Cell Rep. 1:43–55. DOI: 10.1016/j.celrep.2011.11.002

Zill SN (1985). Plasticity and proprioception in insects 1. Responses and cellular properties of individual receptors of the locust metathoracic femoral chordotonal organ. J. Exp. Biol. 116, 435–461. DOI: 10.1242/jeb.116.1.435

Zhou Y, Cameron S, Chang W-T and Rao Y (2012). Control of directional change after mechanical stimulation in drosophila. Molecular Brain 5, 39. DOI: 10.1242/jeb.116.1.435

